# Histone variant H2A.Z is required for plant salt response by regulating gene transcription

**DOI:** 10.1101/2023.07.26.550612

**Authors:** Rongqing Miao, Yue Zhang, Xinxin Liu, Yue Yuan, Wei Zang, Zhiqi Li, Xiufeng Yan, Qiuying Pang, Aiqin Zhang

## Abstract

As a well-conserved histone variant, H2A.Z epigenetically regulates plant growth and development as well as the interaction with environmental factors. However, the role of H2A.Z in response to salt stress remains unclear, and whether nucleosomal H2A.Z occupancy work on the gene responsiveness upon salinity is obscure. Here, we elucidate the involvement of H2A.Z in salt response by analyzing H2A.Z disorder plants with impaired or overloaded H2A.Z deposition. The salt tolerance is dramatically accompanied by H2A.Z deficiency and reacquired in H2A.Z OE lines. H2A.Z disorder changes the expression profiles of large-scale of salt responsive genes, announcing that H2A.Z is required for plant salt response. Genome-wide H2A.Z mapping shows that H2A.Z level is induced by salt condition across promoter, TSS and TES (−1 kb to +1kb), the peaks preferentially enrich at promoter regions near TSS. We further show that H2A.Z deposition within TSS provides a direct role on transcriptional control, which has both repressive and activating effects, while it is found generally H2A.Z enrichment negatively correlate with gene expression level response to salt stress. This study shed light on the H2A.Z function in salt tolerance, highlighting the complex regulatory mechanisms of H2A.Z on transcriptional activity for yielding appropriate responses to particularly environmental stress.

## Introduction

As an adverse environmental stress worldwide, salinization of cultivated land is increasingly imposing severe constraints on plant development and crop yield (Gong et al., 2020). Due to their sessile nature, plants have evolved sophisticated strategies resulting from the complex interactions among robust physiological programs and signaling pathways to cope with saline conditions, and such conditions contribute to the global and rapid activation of genome-wide gene expression (Zhu et al., 2013). Transcriptional regulation of salt stress-responsive genes has been investigated extensively in recent decades, and growing evidence reveals that the epigenetic state of chromatin has profound implications for the regulation of transcriptional activity and leads to appropriate levels of responsiveness (Yang, & Gao, 2018). Epigenetic marks, including DNA methylation, nucleosome remodeling and posttranslational modifications of histones, are well characterized and important for the gene expression or chromosomal structures involved in plant salt tolerance (Ueda, & Seki, 2020).

H2A.Z, a well-conserved variant of canonical histone H2A among eukaryotes, has a significant influence not only on transcriptional regulation but also on genome stability, heterochromatin establishment, cell cycle progression, DNA repair and biotic and abiotic stress responses (Raisner, & Madhani, 2006; Luk et al., 2010; Deal, & Henikoff, 2011). The three H2A.Z-encoding genes *Histone H2A 8* (*HTA8*), *HTA9* and *HTA11* are found in Arabidopsis, and their triple mutants lead to severe defects of development (Yi et al., 2006; Kumar, & Wigge, 2010; Coleman Derr et al., 2012). The chromatin remodeling complex SWI2/SNF2-Related 1 (SWR1) is mainly composed of several components, including *Actin-related protein 6* (*ARP6*), *Photoperiod-independent early flowering 1* (*PIE1*) and *Serrated leaves and early flowering* (*SEF*), which are required for the deposition of H2A.Z into specific loci of chromatin by replacing H2A (Choi et al., 2007). In addition, NRP1 (NAP1-related protein) and NRP2 counteract the activity of the SWR1 complex in chromatin, and their loss causes an overaccumulation of H2A.Z across the genome (Wang et al., 2020). Genome-wide studies of chromatin immunoprecipitation have revealed that H2A.Z is widespread throughout genomes and frequently enriched at the first nucleosome located downstream of the transcriptional start site (TSS) in promoter regions, although it can also be detected in gene bodies at a lower level (Deal, & Henikoff, 2011). The role of H2A.Z in transcriptional regulation depends on its accumulation site, and it displays dual effects on gene expression. The presence of H2A.Z at TSS is associated with transcriptional activation or repression, and when located across gene bodies, it negatively correlates with transcription (Kumar, & Wigge, 2010; Dai et al., 2018; Aslam et al., 2019). Moreover, recent reports proposed that H2A.Z regulates gene expression and may be linked with the posttranslational modification that carries at the +1 nucleosome region and related to the cooperative action with DNA methylation and other histone modifications. H2A.Z monoubiquitination serves as a repressive mark of transcription, while H2A.Z acetylation has a positive effect on transcriptional activation (Gómez-Zambrano et al., 2019; Bieluszewski et al., 2022). Regarding cooperative action, DNA methylation can influence the chromatin structure and affect gene silencing by excluding H2A.Z (Coleman Derr et al., 2012). H2A.Z activates gene expression associated with H3K4me3 modification at promoters, and represses enhancer function by modulating the enrichment level of H3K27me3 and H3K4me3 marks (Dai et al., 2018).

The involvement of H2A.Z in response to drought, temperature, phosphate deficiency and pathogens has revealed the robust regulatory dynamics of H2A.Z under various environmental conditions. Responsiveness to drought stress is associated with H2A.Z enrichment at gene body, H2A.Z occupancy is removed from drought-induced genes, and a decrease in H2A.Z occupancy across gene bodies contributes to the transcriptional activation of drought-responsive genes; in addition, H2A.Z is considered to have a repressive effect and promotes tight regulation under nonstress conditions (Sura et al., 2017). The antagonistic regulation of anthocyanin accumulation between H2A.Z and H3K4me3 is also involved in drought and high light stress responses (Cai et al., 2018). Arabidopsis H2A.Z nucleosomes act as temperature sensors evicted from chromatin under increased temperature, and they regulate transcription changes of ambient temperature-responsive genes and tightly control the key transcription factors PIF4 and HSF1 (Kumar, & Wigge, 2010; Cortijo et al., 2017). H2A.Z has been reported to repress the onset of the Pi starvation response (PSR) by reducing the expression of a number of PSR genes (Smith et al., 2010). Similarly, many immune-responsive genes are repressed by H2A.Z-containing nucleosomes under pathogen attack (Berriri et al., 2016). In addition, PIFs achieve H2A.Z removal through direct interaction with EIN6 ENHANCER (EEN) in response to low far-red (FR) light (Willige et al., 2021). NuA4 controls both the acetylation and deposition of H2A.Z, thereby contributing to the switch between the stress response and autotrophic growth (Bieluszewski et al., 2022).

Acting as an epigenetic mark, H2A.Z evidently influences the gene state and transcription level under certain circumstances, and it likely has designated roles in response to specific signals. To date, the exact function of H2A.Z in salt tolerance remains unknown and its global role in controlling basal transcription and particular role in the gene response to salt stress have not been demonstrated. Here, we describe the biological function of H2A.Z in response to salt stress. We find that H2A.Z deficient mutants exhibit salt sensitivity, and H2A.Z over-accumulated plants gain salt tolerance compare with wild type, suggesting that H2A.Z is required for salt responsiveness. Gene expression profile of these H2A.Z related lines show H2A.Z may negatively regulate the transcription of salt responsive genes. This is partially supporting by the genome wide H2A.Z occupancy pattern at TSS, gene repression asks for a higher H2A.Z level, while induced gene expression do not loss H2A.Z enrichment under salt stress, revealing a dual role of H2A.Z on transcriptional progress upon salt response but generally produce a repressive effect independently wherever H2A.Z locate.

## Materials and methods

### Plant material and stress treatments

*Arabidopsis thaliana* accession Col-0 was used as the wild type (WT) in this study. The T-DNA insertion lines of *hta8* (SALK_095694), *hta9-1* (SALK_054814), *hta11-1* (SALK_017235), *hta9-2* (SALK_151544), *hta11-2* (SALK_031471), *arp6-1* (SAIL_599_G03), *sef-2* (SAIL_1142_C03), *pie1-5* (SALK_096434), *arp6-1sef-2* (CS71694), *nrp1-1* (SALK_117793) and *nrp2-1* (SALK_030348) were obtained from the Arabidopsis Biological Resource Center (ABRC). The *hta9-1hta11-1*, *hta9-2hta11-2* and *nrp1-1nrp2-1* double mutants were generated by genetic crossing. For the over-expression lines of H2A.Z encoding genes and the *HTA8*, *HTA9* and *HTA11* coding regions were amplified and cloned into the gateway vector pDONR222 by BP clonase (Invitrogen, 11789020). Then, these entry clones were recombined into the binary vector pGWB551 by LR clonase (Invitrogen, 11791020). The constructs were verified by restriction enzyme digestion and sequenced to ensure that no errors were produced during amplification and cloning. The Agrobacterium GV3101 strain carrying the 35S::*HTA8*, 35S::*HTA9* and 35S::*HTA11* constructs was used to develop transgenic plants by the floral dipping method. The primers used for genotyping of the mutants are listed in Supplementary Table S7. Sterilized seeds were planted on half Murashige and Skoog (MS) medium plus 1% sucrose with 0.8% agar or in pots filled with uniformly mixed PINDSTRUP substrate, and they were then transferred to a growth chamber after vernalization at 4 ℃ for 3 d and grown at 22 ℃ with 50% relative humidity and 16 h light/8 h dark (150 μM photons m^-2^ s^-1^) photoperiod conditions. Salt stress was applied to three-week-old plants grown in soil treated with 150 mM NaCl for 72 h or directly added to the half MS medium. At least three biological replicates were performed per experiment.

### Phenotypic analysis of plant response to salt stress

For the germination assay, the seeds of Col-0 and mutant lines were harvested from plants growing under the same conditions and stored for one month to maintain the same quality. Surface-sterilized seeds were sown on half MS medium supplemented with or without 150 mM NaCl. Each line contained 80 to 100 seeds for one plate. The germination rates were recorded daily, referring to radicle tip or green cotyledon emergence, and three biological replicates were performed. For root growth, 5-d-old seedlings were transferred into half MS medium containing 150 mM NaCl and then photographed after growth for 7 d. The root lengths were measured using IMAGEJ software. At least 30 seedlings were quantified per line, and each experiment was repeated three times.

### Determination of relative water content (RWC)

Plants were harvested after 150 mM NaCl exposure for 72 h and washed with distilled water, followed by surface drying with filter paper. To determine the RWC, leaves were collected and immediately weighed to quantify the fresh weight (FW). The turgid weight (TW) was obtained after overnight immersion in distilled water, and the dry weight (DW) was obtained after subsequent overnight drying at 50 to 60 ℃. The RWC was calculated based on the following formula: RWC = [(FW − DW) / (TW − DW)] *100%.

### Total chlorophyll content analysis

Plants were washed thoroughly in deionized water. Approximately 0.1 g fresh weight of leaves was incubated in a 5 mL ethanol and acetone mixture (1:2, v/v) for 48 h in the dark. The absorbance of the supernatant was analyzed using a spectrophotometer at 665 nm, 649 nm and 470 nm. Total chlorophyll content was quantified by the sum of Chl a and Chl b. Three biological replicates of each treatment were independently performed.

### Na^+^ and K^+^ content analysis

Plants were washed thoroughly in deionized water. The dry weight of the samples was determined after dehydration in an oven at 80 ℃ for 24 h, and the dry materials were then digested with 2 mL of concentrated nitric acid and perchloric acid at 115 ℃ for 5 h. After centrifugation to remove any debris, the supernatants were analyzed using an Optima 8000 ICP‒OES DV Spectrometer (PerkinElmer, USA) according to the manufacturer’s instructions. Three biological replicates were measured for each sample.

### Histochemical staining for ROS

Three-week-old seedlings were infiltrated with 1% (W/V) 3,3’-diaminobenzidine (DAB) (Sigma-Aldrich) in Tris-HCl (pH 6.5) for H_2_O_2_ staining and 0.5% (W/V) nitrotetrazolium (NBT) (Sigma-Aldrich) blue chloride in HEPES-K buffer (pH 7.0) for superoxide staining. For DAB staining, the samples were covered with 1 mg/mL DAB for 18 h in the dark at 28 ℃ and then transferred to 95% ethanol to decolor. For NBT staining, the samples were first immersed in NBT stain solution until a dark blue color appeared and then decolored in 95% ethanol. Representative photographs were obtained from at least five repeats. To assess the relative labeling intensity and enable a direct comparison between leaves from different genotypes under salt treatment and non-treated conditions, all stained leaves were quantified by using the ImageJ software (Schindelin et al., 2012).

### Histone extraction and western blotting

Histone extraction was performed by slight modifications of the procedure described by Li et al. (2012). Briefly, nuclei were extracted from 1 g of three-week-old seedings and treated for 1 h with 0.4 N H_2_SO_4_ to obtain a histone-enriched extract. The histone protein was precipitated in acetone solution containing 10% (w/v) TCA at −20 ℃ overnight, air dried and resuspended in 4 M urea. Laemmli loading buffer (100 mM Tris-HCl, pH = 6.8, 200 mM DTT, 4% SDS, 20% glycerol, 0.1% bromophenol blue) was added to the histone protein and boiled for 5 min. 10-20 μg of total protein was loaded into 4-12% in the case of total protein or 12% acrylamide gels in the case of histone extracts, then the proteins were transferred into PVDF membrane after separating. The primary antibodies anti-H2A.Z (1:3000, AbClonal, China), histone H3 (1:3000, ab1791, Abcam, England) and secondary antibody goat anti-rabbit IgG (HRP) conjugate (1:3000, ab205718, Abcam, England) were used for immunoblotting. All blot membranes were photographed with the Bio-Rad ChemiDoc XRS after protein detecting by ECL western blotting detection reagents.

### Chromatin immunoprecipitation (ChIP) sequencing

Approximately 4 g leaves harvested from three-week-old seedlings was crosslinked in ice-cold 1× PBS buffer with 1% formaldehyde solution under intermittent vacuum until they became translucent, then quenched by adding glycine to a final concentration of 125 mM and applying vacuum for another 5 min. Plant material was washed in ice-cold milli-Q water twice, pat-dried and ground to a fine powder, resuspended in ice-cold nuclei extraction buffer (10 mM PBS, pH 7.0, 0.1 M NaCl, 10 mM β-mercaptoethanol, 12.8 % 2-methy 2,4-pentanediol and protease inhibitor cocktail (78429, Thermo Scientific, USA) and incubated for 1 h on ice with shaking. The homogenate was filtered through two layers of miracloth (475855, Millipore, USA) and centrifuge at 1000 × g for 20 min at 4 ℃. The pellets were resuspended five times with a total 25 mL nuclei lysis buffer (10 mM PBS, pH 7.0, 0.1 M NaCl, 10 mM β-mercaptoethanol, 12.8 % 2-methy-2,4-pentanediol, 250 μl MgCl_2_, 0.5% Triton X-100 and protease inhibitor cocktail. Chromatin extracts were transferred in the sonic buffer (10 mM PBS, pH 7.0, 0.1 M NaCl, 0.5 % Sarkosyl, 1 mM EDTA, pH 8.0 and protease inhibitor cocktail) and sheared into 200-500 bp fragments with a sonicator (Diagenode, Bioruptor Plus, Belgium). The sonicated chromatin complex was immunoprecipitated by anti-H2A.Z and isolated by binding to Dynabeads Protein A/G (26162, Thermo Scientific, USA). The anti-H2A.Z IP samples and input DNA (which was not immunoprecipitated and serves as a background control) under salt stress were used to construct paired-end library. DNA sequencing was performed on the Illumina sequencing platform (HiSeq4000 PE150). The clean reads were mapped to TAIR10 using BOWTIE2 (v.2.2.3) (http://bowtie-bio.sourceforge.net/bowtie2/index) with default parameters and only uniquely mapped reads were kept. After mapping reads to the reference genome, we used the MACS2 version 2.1.0 (model-based analysis of ChIP-seq) peak finding algorithm to identify regions of IP enrichment over background. A *P* value threshold of enrichment of 0.05 was used for all data sets. Heat maps were generated using the open source tool deepTools.

### RNA isolation and quantitative reverse transcription-(qRT)PCR

Total RNA was extracted from three-week-old plants with Trizol reagent (9109, Takara, Japan). The RNA was treated with gDNA Eraser (RR047A, Takara, Japan) to remove DNA contamination. 2 μg RNA of each sample was used to generate cDNA libraries by NEBNext® UltraTM RNA Library Prep Kit for Illumina® (New England Biolabs). The qRT-PCR was performed using TB Green Premix Ex Taq^TM^ (RR820A, Takara, Japan) with above cDNA as template. Relative expression level of genes was calculated following 2^-ΔΔCt^ method and normalized to the expression level of *ACTIN2*. Three technical replications were conducted for each biological experiment. Mean values represent relative expression level, and error bars standard for deviation (±SD). The primers used in the qRT-PCR analysis are shown in Supplementary Table S5

### RNA sequencing

For RNA-seq library construction, 2 μg of total RNA were used for mRNA purification and cDNA synthesis. The cDNA (100∼200 bp) samples were purified with AMPure XP system (Beckman Coulter, United States). PCR Enriched cDNAs were used to create the final cDNA library and library quality was assessed with the Agilent Bioanalyzer 2100 system (Agilent Technologies). The high-performance sequencing was run on Illumina HiSeq 2500 platform. Clean reads were mapped to the Arabidopsis TAIR10 reference genome and assembly were performed with the software tools HISAT2 and StringTie. Differential expression analysis was performed using DESeq2 package. Genes with an FDR-adjusted fold change ≥2 and *p* ≤ 0.05 were considered to be differentially expressed between two samples. TopGO software was used for gene ontology (GO) functional classification (Alexa, & Rahnefuhrer, 2010; Anders, & Huber, 2010). Three biological replicates were taken for each genotype under control and salt stress conditions.

### ChIP-qPCR

The ChIP-qPCR experiments were performed to validate the changes of H2A.Z enrichment in specific loci of selected genes. DNA enrichment was estimated as the fraction of immunoprecipitated DNA relative to input (% Input). Relative H2A.Z levels were determined as % Input of each region % Input fragment. Sequences for the primers used for ChIP-qPCR are listed in Supplementary Table S7.

### Statistical analysis

Student’s *t* test was used to assess the statistically significant differences between wild type and mutants under salt stress, respectively. Multi-factor ANOVA analysis for genotype-environment interaction (G×E) was performed using Tukey’s multiple comparison test in the RStudio 1.2.1335 (https://www.rstudio.com/).

### Accession numbers

Sequence data in this study can be found in TAIR with the following accession numbers: *HTA8* (AT2G38810), *HTA9* (AT1G52740), *HTA11* (AT3G54560), *ARP6* (AT3G33520), *SEF* (AT5G37055), *PIE1* (AT3G12810), *NRP1* (AT1G74560), *NRP2* (AT1G18800), *WRKY54* (AT2G40750), *KIN1* (AT1G14370), *AKR4C9* (AT2G37770), *MYB75* (AT1G56650), *NPF6.4* (AT3G21670), *OBL1* (AT3G14360), *ALDH7B4* (AT1G54100), *RD22* (AT5G25610), *NAC72* (AT4G27410), *PHD1* (AT3G30775) and *PAL1* (AT2G37040).

## Results

### Plants deficient in H2A.Z show salt sensitivity

To examine the contribution of H2A.Z variants to salt tolerance, we first investigated the roles of H2A.Z-encoding genes in the Arabidopsis response to salinity. The transcripts of *HTA8*, *HTA9* and *HTA11* all showed pronounced induction upon salt stress (Supplementary Fig. S1). Meanwhile, the accumulation of the H2A.Z protein was dramatically promoted under salt stress, indicating that H2A.Z positively responded to salt stress at both the transcript and protein levels (Fig. 1A, B). To further dissect the involvement of H2A.Z in the salt response, T-DNA insertion lines of H2A.Z-encoding genes and knockout mutants were used to analyze the growth phenotype under salt stress. The rosette leaves of the single knockdown mutants *hta8*, and two independent mutant alleles of *hta9* and *hta11* did not show apparent morphological differences compared to the WT under either control or salt stress conditions (Fig. 1C, D, Supplementary Fig. S3 C-E), it is consistent with the previous statement that there was functional redundancy among H2A.Z genes (March-Díaz and Reyes, 2009). Because the triple mutant *h2a.z* show severe dwarfism, to avoid the lethal effect caused by salt stress and better assess the salt response phenotype, therefore we generate the double mutants *hta9hta11*, where the global levels of H2A.Z were drastically reduced compared with that of the WT (Fig. 1C, D, Supplementary Fig. S2A, B, Fig. S3A, B), showed severe wilting and fading after the NaCl treatment and were much more sensitive to salt stress than the WT, supporting by the determination of relative water and chlorophyll content (Fig. 1C, D, Supplementary Fig. S5A, Fig. S3C-E). A similar pattern was observed for root growth, with the root lengths of *hta9-1hta11-1* obviously retarded by salt stress (Fig. 1E, F). Based on the degree of staining with DAB or NBT, the mutant *hta9-1hta11-1* had a higher accumulation of H_2_O_2_ and O_2_ ^-^ than WT under salt stress (Fig. 1G, H). Na^+^ toxicity and salinity-induced K^+^ deficiency are two major constraints in salinized plants, maintaining the balance of Na^+^ and K^+^ is critical for detoxifying excessive Na^+^ ionic damage in salinity-affected plants (Dw et al., 2017). The ratios of K^+^/Na^+^ were significantly decreased in mutants compared with WT under salt conditions due to a higher accumulation of Na^+^ and reduction in K^+^ levels in *hta9-1hta11-1* (Fig. 1I). Delayed germination and green cotyledon emergence were observed in the *hta9-1hta11-1* under the control conditions (Supplementary Fig. S4A, B, Supplementary Table S1), and the difference between these mutants and the WT was enhanced significantly in the NaCl-supplemented medium (Supplementary Fig. S4C, D, Supplementary Table S1), which was validated by the G×E analysis (Supplementary Table S2). Collectively, our observations suggested that plants with H2A.Z deficiency showed higher sensitivity to salt stress.

**Fig. 1.**
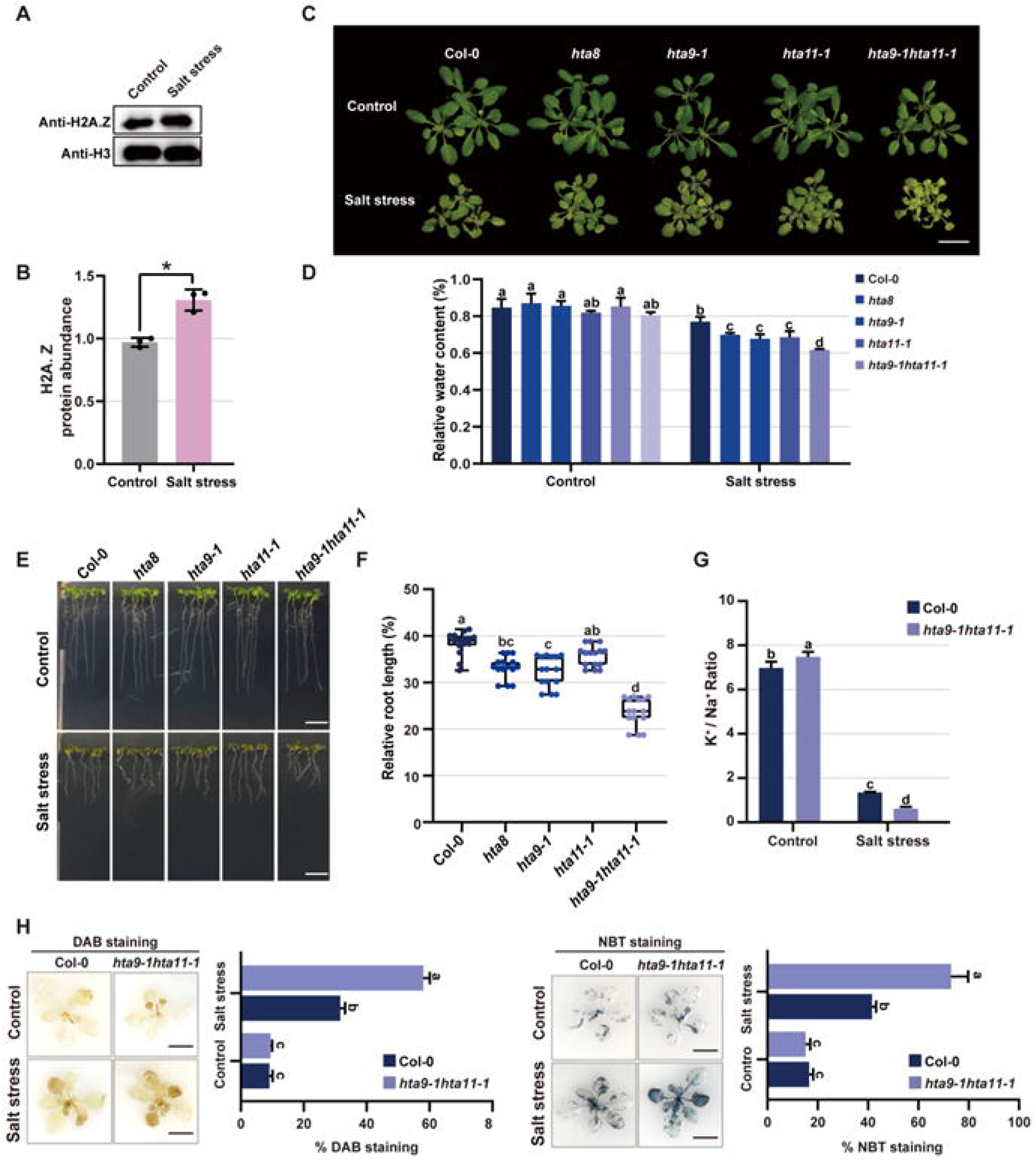
Plants deficient in H2A.Z accumulation showed salt sensitivity. (A) Western blot analysis of histone H3 and variant H2A.Z in Col-0 under control and salt conditions. (B) H2A.Z protein abundance under salt stress detected by using image studio lite from three independent replicates of western blot analysis. Statistical differences between mutants and wild type were analyzed by Student’s *t* test, ** *P* < 0.01. Each bar represents means ± SD, n = 3. (C) Growth phenotype of 3-week-old Col-0 and H2A.Z deficient mutants with or without 150 mM NaCl treatment. Scale bar = 2 cm. (D) The relative water content of Col-0 and H2A.Z deficient mutants under salt stress. (E) The root growth of Col-0 and H2A.Z deficient seedlings on the medium with or without 150 mM NaCl treatment. Scale bar = 1 cm. (F) Quantification of relative root lengths of the seedlings after salt stress. The relative root lengths were calculated by comparing seedlings treated with NaCl to seedlings treated without NaCl. (G) K^+^/Na^+^ ratio in Col-0 and *hta9-1hta11-1* after 150 mM NaCl treatment. (H) DAB and NBT staining of Col-0 and *hta9-1hta11-1* under salt stress. The intensity of DAB and NBT staining was quantified as the stained leaf area relative to the whole leave area (% DAB and % NBT staining). Scale bar = 2 cm. Different letters represent significant difference in each condition (Duncan’s test, *P* < 0.05). Each bar represents means ±SD, n = 15.

### H2A.Z positively regulates Arabidopsis salt tolerance

To further determine the functional role of H2A.Z in salt response, we generated *HTA8*, *HTA9* and *HTA11*-overexpression lines. Immunoblotting showed that the H2A.Z protein over-accumulated in all transgenic lines compared to the WT (Fig. 2A-C). Meanwhile, the double mutant *npr1-1npr2-1* for *NRP1* and *NRP2*, which are negative regulators of H2A.Z abundance of chromatin, and *npr1-1npr2-1* showed overaccumulation of H2A.Z (Supplementary Fig. S2C, D), were also used to dissect the effect of H2A.Z in the salt stress response. The OE lines of H2A.Z-encoding genes displayed healthier leaf morphologies and comparable root lengths as the WT under the control condition, the root lengths were significantly longer than those of the WT under salt stress (Fig. 2A, E, F). The G×E interaction also revealed that *nrp1-1nrp2-1* root development was salt-dependent (Supplementary Table S2). Moreover, the seed germination rate and green cotyledon emergence of the H2A.Z OE lines recovered to that of the WT after the salt treatment (Supplementary Fig. S6, Supplementary Table S1), and the physiological index of the K^+^/Na^+^ ratio, relative water content and ROS accumulation also indicated that plants that over accumulated H2A.Z possessed a salt-tolerant phenotype (Fig. 2D, Supplementary Fig. S7-8), thus revealing that H2A.Z plays a positive role in the regulation of the Arabidopsis response to salt stress.

**Fig. 2.**
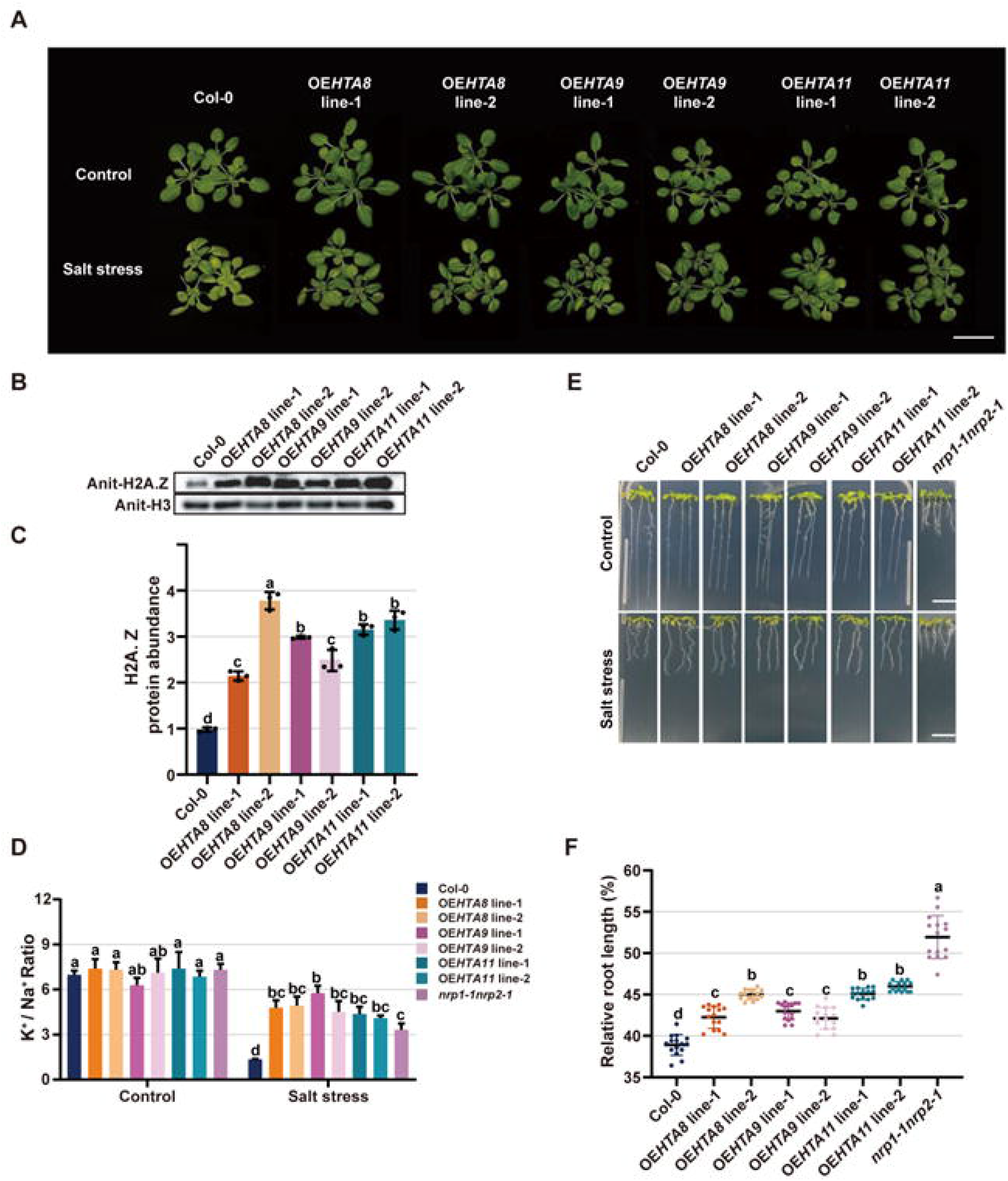
H2A.Z positively regulates salt tolerance. (A) Growth phenotype of 3-week-old Col-0 and H2A.Z overaccumulated seedlings with or without 150 mM NaCl treatment. Scale bar = 2 cm. (B) H2A.Z protein abundance under salt stress detected by using image studio lite from three independent replicates of western blot analysis. (C) H2A.Z protein abundance in Col-0 and H2A.Z overaccumulated seedlings detected by using image studio lite from three independent replicates of western blot analysis. (D) K^+^/Na^+^ ratio in Col-0 and H2A.Z overaccumulated plants after 150 mM NaCl treatment. (E) The root growth of Col-0 and H2A.Z overaccumulated seedlings under control and salt conditions. Scale bar = 1 cm. (F) Quantification of relative root lengths of the seedlings after salt stress. The relative root lengths were calculated by comparing seedlings treated with NaCl to the one without NaCl. Scale bar = 2 cm. Different letters represent significant difference in each condition (Duncan’s test, *P* < 0.05). Each bar represents means ±SD, n = 15.

### H2A.Z deposition is required for the salt stress response

Incorporation of the histone variant H2A.Z into nucleosomes by the SWR1 chromatin remodeling complex is a critical step for performance of H2A.Z functions in eukaryotic gene regulation (Aslam et al., 2019). Therefore, we explored whether the lack of H2A.Z occupancy in nucleosomes affects the plant salt response. H2A.Z mutants deficient in the SWR1 complex components *PIE1*, *ARP6*, and *SEF* were used to examine the overall morphological phenotype and test the salt tolerance. The rosette and root growth results showed that the *arp6-1* and *sef*-2 mutants phenocopied the double mutant *hta9-1hta11-1* in terms of salt sensitivity (Fig. 3A-C, Supplementary Fig. S5B, C). However, *pie1-5* has no significant G×E effect with salt in terms of the root length, indicating that the phenotype of *pie1-5* is potentially influenced by other biological functions of the PIE1 protein (Supplementary Table S2), thus defective lines would display a pleiotropic phenotype. Delayed germination was observed in *arp6-1*, *sef-2* and *arp6-1sef-2* double mutants under salt treatment (Supplementary Fig. S9, Supplementary Table S1). We also found that the decreased ratios of K^+^/Na^+^ and higher accumulation of the H_2_O_2_ and O ^-^ were shown in mutants compared with the WT under salt conditions (Fig. 3D-F). These data suggest that H2A.Z abundance and nucleosome deposition are required for plant salt tolerance.

**Fig. 3.**
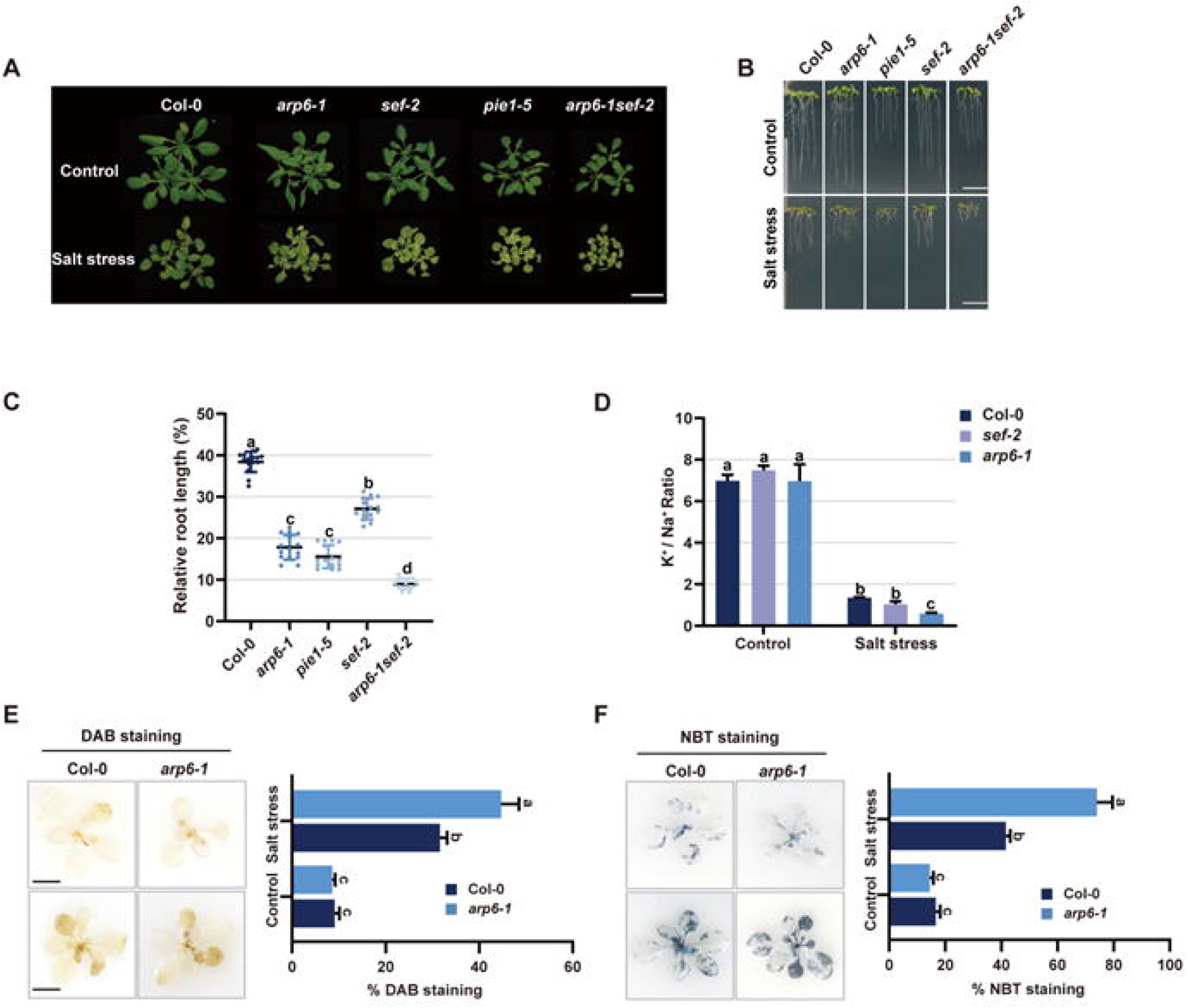
H2A.Z deposition is required for salt stress response. (A) Growth phenotype of 3-week-old Col-0 and H2A.Z depositing defect mutants with or without 150 mM NaCl treatment. Scale bar = 2 cm. (B) The root growth of Col-0 and H2A.Z deficient seedlings under control and salt conditions. Scale bar = 1 cm. (C) Quantification of relative root lengths of the seedlings after salt stress. The relative root lengths were calculated by comparing seedlings treated with NaCl to the one without NaCl. (D) K^+^/Na^+^ ratio in Col-0 and H2A.Z depositing defect plants after 150 mM NaCl treatment. (E)-(F) DAB and NBT staining of Col-0 and arp6-1 under salt stress. Scale bar = 2 cm. Different letters represent significant difference in each condition (Duncan’s test, *P* < 0.05). Different letters represent significant difference in each condition (Duncan’s test, P < 0.05). Each bar represents means ±SD, n = 15.

### Transcription reprogramming of salt responsive genes depends on H2A.Z regulation

Since the H2A.Z variant is distributed throughout the genome and has a robust role in genome-wide transcriptional regulation, we next explored whether H2A.Z generally affects the expression of salt-responsive genes. Transcriptome profiles were generated from H2A.Z-encoding and depositing deficient mutants *hta9-1hta11-1* and *arp6-1* with salt sensitivity and non-significant pleiotropic growth defects, as well as *nrp1-1nrp2-1* with overaccumulation of H2A.Z in chromatin upon salt treatment. The data were analyzed to determine the expression patterns of salt-responsive genes underlying the impaired stress tolerance due to the H2A.Z defect. In the WT background, the differentially expressed genes (DEGs, n=2421) detected under salt stress were referred to as salt-responsive genes and the up (n=1603) and downregulated (n=818) genes were designated salt-induced and salt-repressed genes, respectively (Supplementary Table S3). To examine the impact of H2A.Z deficiency on the transcriptional behavior of salt responsive genes, we compared the gene expression profiles in response to salt stress between H2A.Z depletion mutants and wild type plants. A total of 5664 and 4320 DEGs were identified between *hta9-1hta11-1* and the WT under control condition and salt stress, respectively (Supplementary Fig. S10A, Supplementary Table S4), thus, robust transcriptional changes happened were observed in the plant with less H2A.Z abundance. Moreover, the expression pattern of a large number of genes (n=1633) regulated by *HTA9* and *HTA11* was affected upon salt stress (Supplementary Fig. S10A). A similar phenomenon was observed in the *arp6-1* and *nrp1-1nrp2-1* mutants (Supplementary Fig. S10B, C). Further analysis found that a large set of genes regulated by H2A.Z were salt-responsive genes, with 1106 genes modulated by H2A.Z depletion identified from *hta9-1hta11-1* and 682 genes modulated by H2A.Z overaccumulation identified from the *nrp1-1nrp2-1* background (Fig. 4A). Moreover, almost 50% of the ARP6-regulated genes (486 out of 1015 genes) were shared with the DEGs associated with salt response (Fig. 4A). Therefore, the transcriptome analysis showed a significant overlap between the genes regulated by H2A.Z and the DEGs in response to salt stress, implying that H2A.Z is involved in the transcriptional regulation of salt responsive genes.

**Fig. 4.**
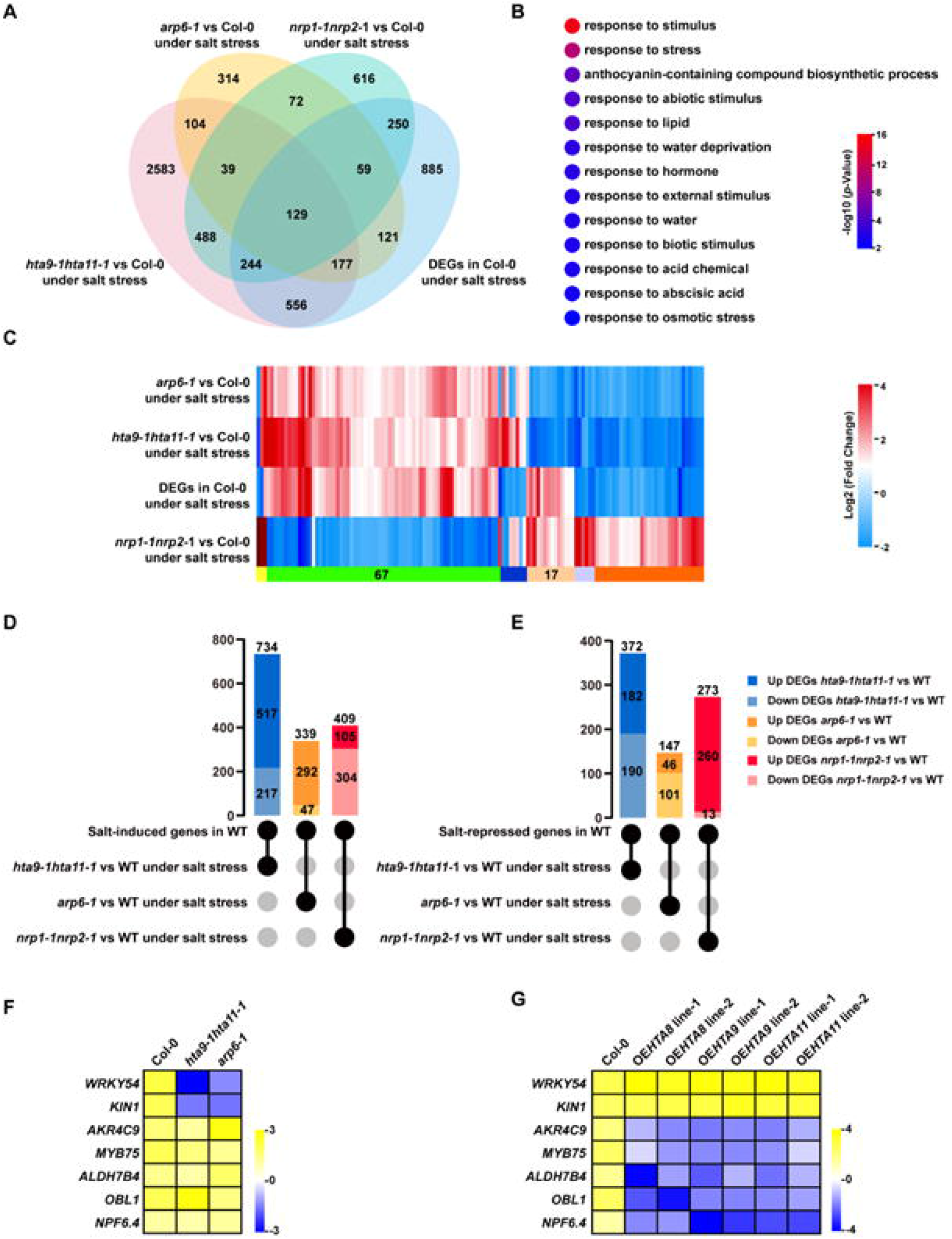
Transcription reprogramming of salt responsive genes depends on H2A.Z regulation. (A) Venn diagram shown the number of overlapping genes between differentially expressed genes (DEGs) in Col-0 and *arp6-1*, *hta9-1hta11-1*, *nrp1-1nrp2-1* under salt stress. (B) GO term enrichment of the overlapping genes between Col-0 and H2A.Z related mutants. (C) Heatmap represents the expression pattern of the overlapping genes detected in Col-0 and H2A.Z related mutants. (D)-(E) The matrix diagram shows different groups of comparison among these gene sets obtained in Col-0 and H2A.Z mutant background; the vertical bar chart indicates the gene number for a given comparison, black circles and gray circles indicate gene sets included and excluded from a certain comparison. (F)-(G) Expression levels of several salt-responsive genes in Col-0, *hta9-1hta11-1*, *arp6-1* and H2A.Z OE lines. Yellow represents upregulation, blue represents downregulation.

Then, we examined whether the salt respond DEGs were shared in the H2A.Z genetic material. An overlapping gene analysis showed that 129 salt-responsive genes were coregulated by *HTA9/11*, *ARP6* and *NRP1/2*, and their expression was highly correlated with H2A.Z enrichment (Fig. 4A). A Gene Ontology (GO) enrichment analysis confirmed that these coregulated genes were preferentially associated with response to stimulus and response to stress (Fig. 4B). A heatmap generated by hierarchical clustering analyses revealed that most (67 out of 129) of the H2A.Z-regulated salt-responsive genes were expressed with the same pattern in H2A.Z-deficient plants but showed opposite trends in H2A.Z over-accumulated plants (Fig. 4C). This result is consistent with previous findings that H2A.Z has a repressive role in global transcription. However, a small set of genes (17 out of 129) exhibited the reverse phenomenon, which was likely because of the positive involvement of H2A.Z in the regulation of expression of these genes (Fig. 4C).

To better understand the regulatory pattern of H2A.Z on salt-responsive genes, we analyzed the expression changes of salt-induced and salt-repressed genes in H2A.Z-deficient and H2A.Z-enriched plants. For salt-induced genes, a large portion of shared DEGs detected in *hta9-1hta11-1* (70%, 517 out of 734) and *arp6-1* (86%, 292 out of 339), were upregulated compared to the WT under salt stress, especially in the *arp6-1* background, and only 47 genes lost their induced expression at a lower level of H2A.Z enrichment under salt conditions. However, most of these genes were downregulated in *nrp1-1nrp2-1* (75%, 304 out of 409) (Fig. 4D). A similar tendency was observed in the set of salt-repressed genes; the repression of many genes in the WT was maintained or enhanced in *hta9-1hta11-1* and *arp6-1* mutants, whereas these genes were promoted in the *nrp1-1nrp2-1* plants with excess H2A.Z (Fig. 4E). Moreover, there was also quite a part of salt-induced or repressed genes failed to be regulated upon H2A.Z deficiency. qRT‒PCR tests for several well-known salt-related genes confirmed the expression change in the reverse context of H2A.Z accumulation (Fig. 4F, G). Stress-responsive markers (eg. *NPF6.4*, *MYB75*, *OBL1*, *ALDH7B4* and *AKR4C9*) were upregulated in *hta9-1hta11-1*, *arp6-1* and the WT, but showed the opposite trend in *HTA8*/*9*/*11* OE lines (Fig. 4F, G). These results suggested a dual role of H2A.Z for triggering or suppressing the expression of salt-responsive genes, and transcriptional changes showed that H2A.Z negatively influences gene expression under salt stress.

### H2A.Z enrichment level induced by salt stress

Given our findings that H2A.Z modulates salt responsiveness, the disruption of genes encoding or depositing H2A.Z led to the mis-regulated expression of general salt-responsive genes. We hypothesized that salt signals might affect genomic H2A.Z occupancy, which then mediates transcriptional processes and salt adaptation. To investigate the change in H2A.Z enrichment upon salt stress, we performed chromatin immunoprecipitation followed by genome-wide sequencing (ChIP-seq) based on H2A.Z-bound DNA. This gave the identification of 32,894 H2A.Z peaks located at 21,196 genes in Col-0 under non-stressed conditions and 33,263 peaks at 21,171 genes under salt stress, shown significantly higher peak occupancies of H2A.Z across whole genome upon salt exposure, 1629 peaks were induced by salt with non-appearance under control (Supplementary Table S8, S9, Supplementary Fig. S11A). More than 80% of the peaks were found in promoter regions close to the transcription start site (TSS), which is consistent with the previous findings that H2A.Z is preferentially enriched at promoters. The peaks were also compatible with published histone H2A.Z ChIP-seq datasets and exhibited approximately 80% overlap (Dai et al., 2018; Gómez-Zambrano et al., 2018). In addition, a small portion of H2A.Z peaks (6%) accumulated at the gene body region predominantly at exons other than the first (Supplementary Fig. S11C). Then, we examined the enrichment level of H2A.Z over genes 1 kb upstream of 1 kb up stream of transcription start sites (TSS) and 1 kb downstream of the transcription ending sites (TES) in the WT. A metagene analysis revealed a substantial increase in H2A.Z abundance under salt stress conditions compared to nonstress conditions around TSS and gene body regions but not 1 kb up stream of transcription start sites (TSS) (Fig. 5A, B), suggesting that salt signals induced H2A.Z enrichment at the genomic level.

**Fig. 5.**
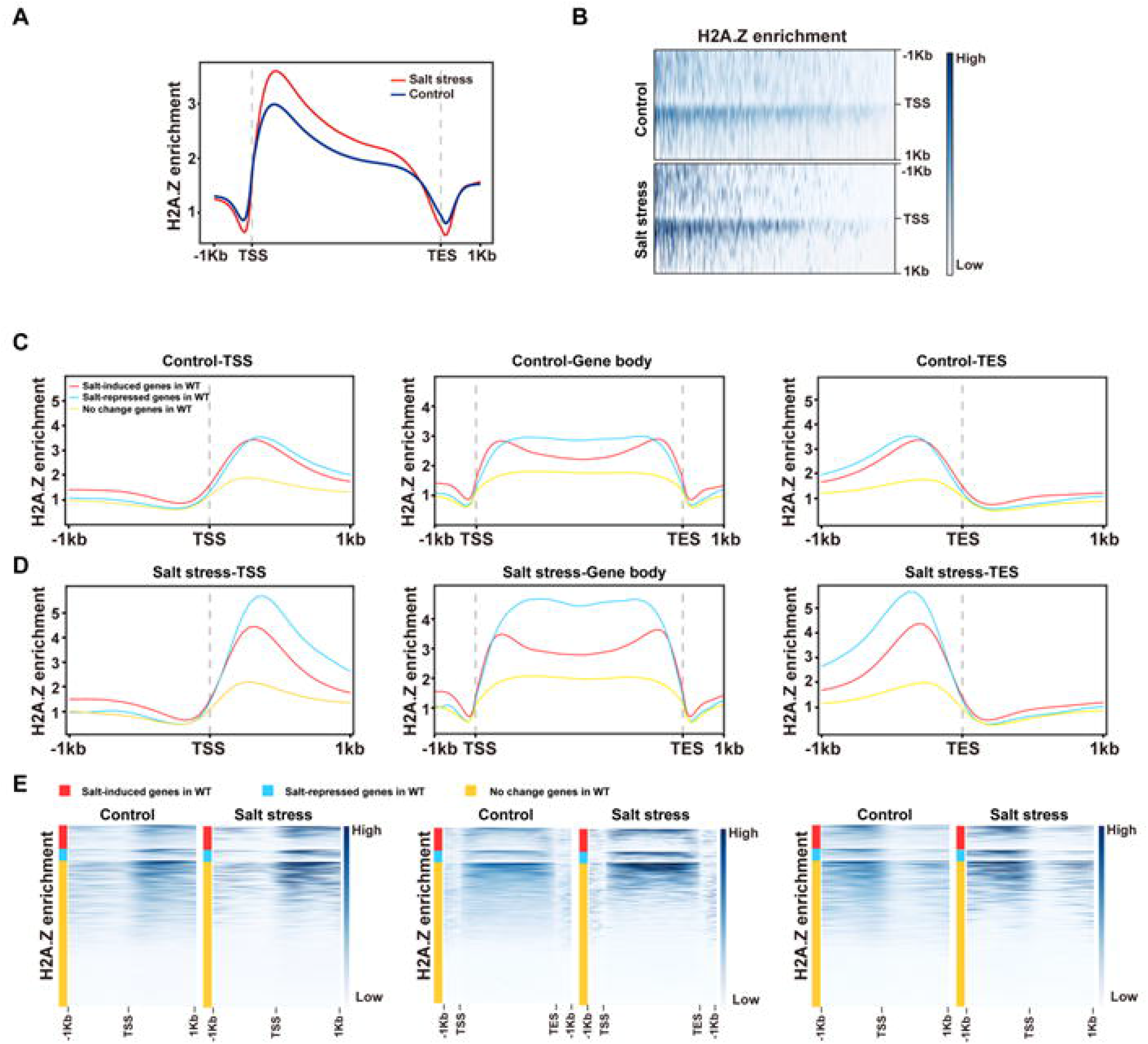
The level of H2A.Z enrichment induced by salt stress. (A) Metagene plots shown H2A.Z levels over all genes including the regions −1 kb of TSS (transcription start site), +1 kb of TES (transcription ending site) and gene body under control and salt stress. (B) Heatmap shown the distribution of H2A.Z across genome under control and salt conditions. (C)-(D) H2A.Z enrichment of salt-induced (red line) and salt-repressed (blue line) genes relative to TSS, TES, and gene body region in comparison with “no change genes” (yellow line). (C) Control. (D) Salt stress. (E) Heatmap visualization of H2A.Z enrichment at TSS/TES/gene body represented in genes under control and salt stress.

To determine the correlation between the expression of salt-responsive genes and H2A.Z occupancy, the H2A.Z accumulation pattern was checked in three sets of genes, two of which were defined based on the RNA-seq analysis as “salt-induced genes” and “salt-repressed genes”. The genes with no expression change under salt stress were grouped as “no change genes”. H2A.Z enrichment was observed at three regions independently, near the TSS, TES and gene body. The metagene plot and heatmap showed greater H2A.Z abundance at all gene regions of salt-induced and repressed genes compared to the no change genes (Fig. 5C-E), thus revealing that the transcriptional activity of the gene response to salt stress was dynamically modulated by H2A.Z occupancy. Furthermore, the level of H2A.Z abundance in salt-repressed genes was much higher than that in salt-induced genes, which indicates the negative role of H2A.Z in regulating gene expression for salt responsiveness. Altogether, these results confirm that H2A.Z deposition across gene bodies is required for the transcriptional response to salt stress.

### H2A.Z deposition level is associated with the expression pattern of salt responsive genes

The global transcription of salt-responsive genes in the H2A.Z-deficient mutants *hta9-1hta11-1* and *arp6-1*, as well as in the H2A.Z overaccumulation line *nrp1-1nrp2-1* was altered, which indicated that nucleosomal H2A.Z deposition affects gene transcription involved in the salt response (Supplementary Fig. S11D-F). The expression change pattern of these genes under H2A.Z absence or excess background revealed a generally negative role of H2A.Z on gene transcriptional level. To further determine how H2A.Z occupancy is associated with gene expression patterns under salt stress, we compared the H2A.Z enrichment level of salt-induced and repressed genes with the misregulated salt-related genes in the mutants *hta9-1hta11-1*, *arp6-1* and *nrp1-1nrp2-1*, and the genes were separated into hyperactive and hypoactive sets, which corresponded to the up and downregulated genes in mutants, respectively. The ChIP-seq profiles showed that the majority of H2A.Z peaks were distributed in promoter regions around TSS (Supplementary Fig. S11C); therefore, we focused on the deposition pattern of H2A.Z near the TSS (−1 kb to +1 kb) region to analyze the correlation between the gene expression level and H2A.Z enriched abundance.

For salt-induced genes, most salt-induced genes were hyperactive in the mutants *hta9-1hta11-1* and *arp6-1* but hypoactive in *nrp1-1nrp2-1*, as demonstrated by RNA-seq data (Fig. 4D, E). There were no differences in H2A.Z distribution near the +1 kb TSS of these genes under the control conditions, although the hyperactive genes in *hta9-1hta11-1* and *arp6-1* showed more H2A.Z enrichment than the hypoactive genes in *nrp1-1nrp2-1* upon salt stress followed by the H2A.Z distribution pattern in WT (Fig. 6A-C). This finding showed that genes were hyperactive under a deficiency in H2A.Z encoding or deposition induced by the salt treatment, thus revealing that H2A.Z negatively mediated the gene transcriptional level. The hypoactive genes detected in *hta9-1hta11-1* and *arp6-1* exhibited relatively lower H2A.Z levels in comparison with the salt-induced genes (Fig. 6B, C). Moreover, hyperactive genes in *nrp1-1nrp2-1* showed marginally higher levels of H2A.Z at +1 kb TSS under both control and salt conditions, which indicated enhanced H2A.Z occupancy under salt stress (Fig. 6B, C). These results indicated that the observed alterations in gene expression are a direct result of changes in H2A.Z levels. Thus, the transcriptional regulation of gene responses to salt required certain level of H2A.Z accumulation at +1 kb TSS. A very similar mode was detected in salt-repressed genes, which were likely hyperactive in nrp1-1nrp2-1 (Fig. 6D-F). We found higher H2A.Z enrichment across the hyperactive genes. Moreover, the hypoactive genes in *hta9-1hta11-1* and *arp6-1* had less H2A.Z than *nrp1-1nrp2-1* at the TSS under the control and stress conditions, and the discrepancy was enlarged by salt stress. These hypoactive genes also did not lose H2AZ upon transcriptional repression (Fig. 6D-F). These findings confirmed that a direct effect of H2A.Z occupancy level on transcription, and somehow H2A.Z produce both active and repressive effect on the expression of salt-responsive genes. Then we randomly selected five salt-responsive genes putatively targeted by H2A.Z and performed ChIP-qPCR analysis to assess H2A.Z occupancy at specific site. H2A.Z significantly enriched at TSS of these five genes under both control and stress conditions in the WT, *nrp1-1nrp2-1* and OE*HTA9* line-1, but extremely weak signal of H2A.Z shown in *hta9-1hta11-1* and *arp6-1* because of the deficiency in H2A.Z deposition (Fig. 6H). Comparing to control condition, ChIP-qPCR analysis showed a decrease enrichment of H2A.Z near TSS of *RD22*, *NAC72* and *WRKY54*, but an increase in *PHD1* and *PAL1* under salt stress in Col-0, *nrp1-1nrp2-1* and OE*HTA9* line-1 which was consistent with the observation in ChIP analysis as IGV map displayed (Fig. 6G, H). The gene expression analysis of these H2A.Z targets revealed a negative correlation between the H2A.Z occupancy near TSS with their expression level upon salt treatment (Fig. 6H), indicating the expression change of these genes response to salt was regulated by H2A.Z enrichment.

**Fig. 6.**
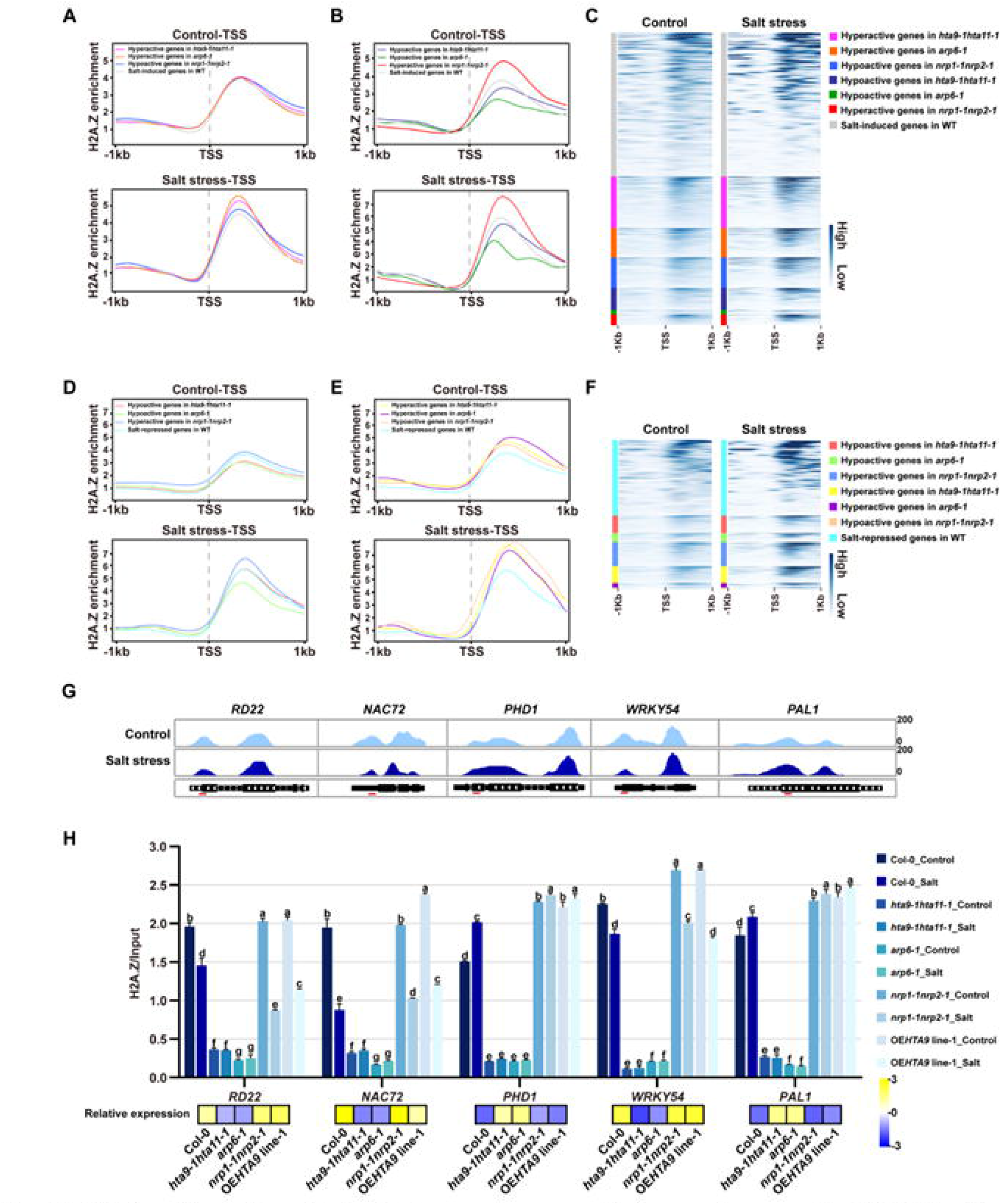
H2A.Z deposition level associated with the expression pattern of salt responsive genes. (A)-(B) H2A.Z enrichment level of WT in the hyper or hypoactive genes in *hta9-1hta11-1* and *arp6-1* and hyper or hypoactive genes in *nrp1-1nrp2-1* compared with salt-induced genes under control and salt stress. (C) Heatmap visualization of H2A.Z enrichment at TSS of the genes represented in (A) and (B) under control and salt stress. (D)-(E) H2A.Z enrichment level of in the hyper or hypoactive genes in *hta9-1hta11-1* and *arp6-1* and hyper or hypoactive genes in *nrp1-1nrp2-1* compared with salt-repressed genes under control and salt stress. (F) Heatmap visualization of H2A.Z enrichment at TSS of the genes represented in (D) and (E) under control and salt stress. (G) Browser views of H2A.Z levels in Col-0 at specific locus of the selected genes, and schematic representation of each gene with red lines indicating the position of amplified regions. (H) ChIP qRT-PCR analysis of H2A.Z enrichment at target loci in the selected genes in Col-0, *hta9-1hta11, arp6-1*, *nrp1-1nrp2-1* and OE*HTA9* line-1 under control and salt stress. Anti-H2A.Z ChIP was normalized to input. Yellow and blue represents the expression levels of *RD22*, *NAC72*, *PHD2*, *WRKY54* and *PAL1* in Col-0, *hta9-1hta11, arp6-1*, *nrp1-1nrp2-1* and OE*HTA9* line-1 under salt stress. Different letters represent significant difference in each condition (Duncan’s test, *P* < 0.05).

## Discussion

To withstand extreme environmental conditions, genetic factors, especially epigenetic changes, play a decisive role in stress responsiveness via transcriptional regulation. In recent decades, considerable evidence has shown that the histone variant H2A.Z functions as a crucial epigenetic mark to maintain genome stability and chromatin accessibility and has regulatory effects on multiple biological processes and environmental stimulus responses (Kumar, & Wigge, 2010; Billon, & Cote, 2012). However, whether H2A.Z contributes to salt tolerance remains unclear. In this work, we assessed the salt tolerance of plants with defects or excesses in H2A.Z deposition. The H2A.Z-encoding and depositing-deficient mutants displayed hypersensitivity to salt stress, resulting in an imbalance of ROS and K^+^/Na^+^, while the *HTA8/9/11*-OE lines and *nrp1-1nrp2-1* mutants displayed a salt-tolerant phenotype (Fig. 1-3), suggesting that H2A.Z is required for the salt response. Because of the importance of appropriate nucleosome H2A.Z occupancy in plant development, a significant growth defect was observed in *pie1-5*, *arp6-1sef-2*, *h2a.z* and *nrp1-1nrp2-1* mutants, in which the impact of the mutation on the salt-responsive phenotype was not observed. Genotype and environment interaction analyses based on root growth preliminarily confirmed a molecular link between H2A.Z and salt tolerance. The conjoint analysis of ChIP-seq and RNA-seq enabled us to determine how H2A.Z mediates transcriptional changes involved in the proper response. H2A.Z enrichment across gene bodies was induced after salt stress, and the changes were negative correlated with the expression level of most salt-related genes. Moreover, their occupancy mode indicated that H2A.Z levels determine the gene responsiveness to salt stress based on a dual role.

### H2A. Z is key for the transcriptional activity of salt-responsive genes

The role of H2A.Z in transcriptional control has been well reported, although the exact mechanisms of H2A.Z occupancy in relation to transcriptional regulation have not been clarified. Conflicting evidence indicates that H2A.Z has both an activating and a repressive effect on transcription (Giaimo et al., 2019; Lewis et al., 2021). Several studies on the Arabidopsis response to stimuli support the repressive role of H2A.Z in modulating gene expression (March-Díaz et al., 2008; Kumar, & Wigge, 2010; Smith et al., 2010). As our investigation showed, a considerable number of salt-responsive genes were deregulated in H2A.Z-deficient or overaccumulation mutants (Fig. 4A, B), and most of these genes changed their expression patterns upon aberrant H2A.Z deposition. The genes induced by salt stress generally present higher expression in H2A.Z-deficient plants but decreased abundance in *nrp1-1nrp2-1* plants with excess H2A.Z accumulation. In contrast, salt-repressed genes exhibited the opposite expression pattern in *arp6-1* and *nrp1-1nrp2-1* (Fig. 4C), which is consistent with the role of H2A.Z in negatively regulating gene expression programs. These results may be inconsistent with the observation of a salt-sensitive phenotype in the H2A.Z-deficient mutant, which implies a positive role for H2A.Z. A similar finding was reported in an analysis of the response to drought stress (Sura et al., 2017). However, some salt-responsive genes are misregulated when the synthesis of H2A.Z protein is insufficient. In *hta9-1hta11-1*, approximately 30% of salt-induced genes lost their expression (Fig. 4E, F), thus indicating that H2A.Z also acts as an activator of environmentally influenced gene expression. Thus, we could conclude that H2A.Z performs a two-way function in the transcriptional regulation response to salt stress and generally exerts a repressive effect. It has been proposed that H2A.Z represses basal transcription but is required for full gene responsiveness at nucleosomes (Dai et al., 2018), thus revealing the regulatory complexities of H2A.Z responses to environmental conditions. The methods by which these two regulatory roles coordinate transcriptional control across the genome and the specificity or preference for H2A.Z distribution at distinct genes or gene sets with regard to the varying contributions to transcription must be further investigated.

### The genomic occupancy of H2A.Z dually correlated with gene expression upon salt stress

Previous studies showed that the H2A.Z peak is dominantly enriched in the first (+1) nucleosome immediately after TSS in eukaryotes (Raisner et al., 2005; Barski et al., 2007; Daniel et al., 2008; Gómez-Zambrano et al., 2018). This is consistent with our investigation, which showed that the genome-wide scale of H2A.Z peaks were mainly distributed in the promoter region near the TSS while a small portion of the peaks located in the exons of the gene body. After salt stress, we observed a significant induction of H2A.Z peaks across the promoter and gene body, thus reflecting the dynamic change in H2A.Z occupancy in the genome upon salt treatment, which is supported by the overall accumulation of H2A.Z according to immunoblot detection (Fig. 5A, Fig. 1A, B).

To study the correlation of transcriptional regulation with locus-specific H2A.Z enrichment level under stress conditions, the genomic H2A.Z occupancy was checked through salt-responsive genes and genes with altered expression in H2A.Z-related mutants under both control and salt conditions. This analysis revealed a dual correlation between H2A.Z enrichment and transcriptional abundance. Regardless of whether the gene is induced or repressed after salt stress, higher H2A.Z enrichment was found across whole gene regions, suggesting that H2A.Z enrichment associated with both transcriptional repression and activation occurs (Coleman Derr et al., 2012; Sura et al., 2017). While the H2A.Z level in salt-repressed genes was higher than that in induced genes (Fig. 5), a generally repressive role of H2A.Z was observed. A negative correlation was observed between H2A.Z at the gene body and transcription levels, and transcriptional activation upon drought stress depends on decreases in the occupancy of H2A.Z across gene bodies H2A.Z at the +1 nucleosome of some genes led to the positive regulation of transcriptional activity (Sura et al., 2017). This was discrepancy with this study, we found that H2A.Z enrichment around TSS including +1 nucleosome peak has two-way influence on gene expression (Fig. 6). It may reveal that H2A.Z work with distinct model on transcriptional reprogramming upon different environmental factors.

### The potentially regulatory mode of H2A.Z on salt responsiveness

A relative lower H2A.Z accumulation was observed at TSS of the hyperactive or hypoactive genes in H2A.Z mutants, also salt-induced and repressed genes under control conditions (Fig. 6A-F), indicating that nuclesomal H2A.Z occupancy was required for basal transcription. The hyperactive genes due to H2A.Z defect, had higher H2A.Z enrichment than hypoactive genes due to excess H2A.Z under salt stress based on the H2A.Z deposition profiles in WT background (Fig. 6A-C), and these hyperactive genes caused by excess H2A.Z shown more abundant H2A.Z level, these observations again support a repressive effect of H2A.Z on the expression of salt responsive genes. Consistent with what we found, it has been proposed the Arabidopsis H2A.Z at +1 nucleosome near TSS inhibit transcription by a closed nucleosome structure and decreased local chromatin accessibility (Dai et al., 2018). On the other hand, the transcriptional activation of some genes also needs a certain level of H2A.Z enrichment, which supports a dual role of H2A.Z in transcriptional control.

There are at least two possible explanations for the regulatory role of H2A.Z in response to salt stress. First, salt induced the overall H2A.Z level across genome, the increase of H2A.Z occupancy may affect the coordination with histone modification markers like methylation and acetylation and change chromatin accessibility dynamics. The cooperation of H2A.Z with active or repressive histone modification markers would generate a direct effect on transcription by providing dual effects of H2A.Z on gene accessibility, which could mediate the binding of positive and negative regulators with specific genes. It has been reported that environmental responsive genes are associated with highly accessible chromatin in Arabidopsis, H2A.Z associated with H3K4me3 at promoters to activate gene expression, and repressed enhancer function by promoting the repressive H3K27me3 marks and inhibiting H3K4me3. An alternative explanation is that H2A.Z dynamically modulate RNA polymerase II (RNAPII) initiation and elongation, RNAPII presents at conserved +1 H2A.Z nucleosome, this could impact the preinitiation complex orientation and transcriptional elongation, several studies stated that H2A.Z occupied at +1 nucleosomes may maintain Pol II in a paused state, which is important for the proper transcription (Kumar, & Wigge, 2010; Nock et al., 2012; Rhee, & Pugh, 2012; Mylonas et al., 2021). Taken together, our analyses support an essential role of H2A.Z in regulating salt tolerance by dually modulating transcriptional activity. We found H2A.Z enrichment was significantly induced by salt, most of the peaks located around +1 nucleosomes after TSS. The nucleosome occupancy of H2A.Z provided dual effects on transcriptional control of salt responsive genes. Further precise work on the regulatory complexities of H2A.Z response to salt is needed to investigate, of how H2A.Z gives tight regulation on transcriptional activity with both active and repressive role, and for the dual effect, what the preference of H2A.Z targets to different genes in whole genome. In addition, it will be worth dissecting the association of H2A.Z with other epigenetic players like histone modifications involved in transcriptional control, then offer a proper responsiveness upon salt stress.

## Author contributions

Aiqin Zhang, Qiuying Pang and Xiufeng Yan designed the experiments and supervised the project. Rongqing Miao, Yue Zhang, Xinxin Liu, Yue Yuan, Wei Zang and Zhiqi Li performed the experiments and integrated the data. Rongqing Miao and Aiqin Zhang wrote the manuscript with contributions from other authors.

## Data availability

Data are available from the corresponding authors, Aiqin Zhang and Qiuying Pang, upon request.

## Supplementary data

Fig. S1. The expression analysis of H2A.Z encoding and depositing genes under salt stress.

Fig. S2. Global level of H2A.Z protein in *hta9-1hta11-1* and *nrp1-1nrp2-1* mutants.

Fig. S3. The growth phenotype analysis of Col-0, *hta9-2*, *hta11-2* and *hta9-2hta11-2* under salt stress.

Fig. S4. Radicle tip and green cotyledon emergence rate in the wild type and H2A.Z deficient mutants.

Fig. S5. Total chlorophyll content and the relative water content of Col-0, H2A.Z deficient mutants, H2A.Z depositing defect mutants and H2A.Z over-accumulated seedlings under salt stress.

Fig. S6. Radicle tip and green cotyledon emergence rate in the wild type and H2A.Z overaccumulated seedlings.

Fig. S7. K^+^ and Na^+^ contents in Col-0 and H2A.Z related plants under salt stress.

Fig. S8. DAB and NBT staining for rosette leaves of wild type and H2A.Z over-accumulated plants under 150 mM NaCl treatment.

Fig. S9. Radicle tip and green cotyledon emergence rate in the wild type and H2A.Z depositing defect mutants.

Fig. S10. Venn diagram showing overlaps of differentially expressed genes (DEG)s in *hta9-1hta11-1*, *arp6-1* and *nrp1-1nrp2-1* at control and salt treatment conditions.

Fig. S11. H2A.Z enrichment affects global gene expression under salt stress conditions.

Table S1. Germination assay in wild type, H2A.Z disorder mutants and H2A.Z overaccumulated plants under salt stress.

Table S2. Genotype by environment analysis for H2A.Z related genes in plant growth under salt stress.

Table S3. Differentially expressed genes detected in Col-0 response to salt stress.

Table S4. Differentially expressed genes detected in *hta9-1hta11-1* under control and salt stress.

Table S5. Differentially expressed genes detected in *arp6-1* under control and salt stress.

Table S6. Differentially expressed genes detected in *nrp1-1nrp2-1* under control and salt stress.

Table S7. Primers used in this study.

Table S8. Genome-wide distribution of H2A.Z peaks in Col-0 under control and salt stress.

Table S9. H2A.Z occupied genes under control and salt stress.

